# Vertical Sleeve Gastrectomy inhibits 11βHSD1 and subsequently reduces IL6 secretion in Mice and Humans: A Shared Anti-Inflammatory Mechanism

**DOI:** 10.64898/2026.05.12.724611

**Authors:** Shiyi Liang, Suhanya Samarasinghe, Brett Johnson, Isabella Doria Durazzo, Wenya Wang, Hio Lam Phoebe Tsou, Antonio Riva, Alexander D. Miras, Elina Akalestou

## Abstract

**Background:** Vertical sleeve gastrectomy (VSG) improves glycaemic control in type 2 diabetes (T2D) through mechanisms that extend beyond weight loss. The interaction between glucocorticoid metabolism and inflammation in this context remains unclear.

**Methods:** We investigated the role of 11β-hydroxysteroid dehydrogenase type 1 (11βHSD1) in mediating the metabolic effects of VSG in humans and mice. Subcutaneous adipose tissue biopsies were collected before and 6 months after VSG. Parallel studies were conducted in lean and high-fat diet-fed mice undergoing VSG or sham surgery, alongside 11βHSD1 knockout models. Glucose tolerance and expression of 11βHSD1 and interleukin-6 (IL6) were assessed. Mechanistic interactions were examined in IL6-treated human hepatocytes.

**Results:** VSG reduced 11βHSD1 and IL6 expression in human adipose tissue and improved insulin resistance. In lean mice, VSG improved glucose tolerance and downregulated both markers independently of weight loss. 11βHSD1 knockout mice exhibited improved glucose tolerance despite increased adiposity, partially recapitulating the VSG phenotype. Both interventions reduced circulating and tissue IL6 levels. IL6 stimulation increased HSD11B1 expression in hepatocytes.

**Conclusions:** 11βHSD1 links glucocorticoid metabolism, inflammation, and glucose homeostasis following VSG. Targeting this pathway may offer a strategy to replicate key metabolic benefits of metabolic bariatric surgery.

## Introduction

While numerous pharmacological interventions have been developed for diabetes, more recent strategies have increasingly focused on weight loss and its long-term metabolic benefits (1, 2). Metabolic surgery, including vertical sleeve gastrectomy (VSG), was initially developed as a weight-loss intervention (3) but has since emerged as an effective treatment for T2D, improving glycaemic control, enhancing insulin sensitivity, and reducing cardiovascular risk (4-6). The mechanisms underlying these benefits have been widely investigated, particularly to replicate the metabolic effects of metabolic bariatric surgery through non-surgical approaches.

Several mechanisms have been proposed; both weight loss dependent and independent. These include enhanced secretion and action of gastrointestinal hormones such as glucagon-like peptide-1 (GLP-1) (7) as well as reductions in systemic inflammation (8, 9). Pro-inflammatory cytokines, including interleukin-6 (IL6), are significantly reduced post-surgery in humans (8, 9). Additional reported adaptations include improved pancreatic islet cell connectivity (10), reduced risk of cardiovascular diseases (11), and improved kidney function (12).

Cortisol, a glucocorticoid produced by the adrenal cortex under the regulation of the hypothalamic–pituitary–adrenal (HPA) axis (13), plays a key role in metabolic homeostasis. Its local activation is mediated by 11β-hydroxysteroid dehydrogenase type 1 (11βHSD1), which converts inactive cortisone to cortisol in an NADPH-dependent manner (14) (15). 11βHSD1 is widely expressed, particularly in liver and adipose tissue, and dysregulation of this pathway has been implicated in obesity and T2D, contributing to insulin resistance and impaired glucose control (16-19). Notably, increased 11βHSD1 activity in adipose tissue has been observed in obesity (18, 20), while tissue-specific overexpression studies in mice demonstrate its causal role in metabolic dysfunction (18) (20).

Emerging evidence also highlights a bidirectional relationship between glucocorticoid metabolism and inflammation. While glucocorticoids are classically anti-inflammatory, stress-induced increases in corticosterone have been shown to exacerbate inflammatory responses in mice (21). Human studies further suggest that inflammation can influence cortisol-mediated glucose regulation, as demonstrated in the KORA-Age study (22), while cytokines such as IL-6 activate the HPA axis and increase circulating cortisol levels (23).

The interaction between VSG and glucocorticoid metabolism remains under-explored. Our group previously demonstrated that VSG fine-tunes cortisol secretion in obese mice through downregulation of 11βHSD1 in liver and adipose tissue (24). One potential mechanism underlying this effect may involve the interplay between glucocorticoid regulation and inflammation (22). However, the extent to which 11βHSD1 contributes to the glucose-lowering effects of VSG remains unclear. In this study, we aimed to define the role of 11βHSD1 in mediating the metabolic benefits of VSG in both human and rodent models.

## Methods

Humans - The BAMBINI randomised controlled clinical trial (ISRCTN16668711) was a multicentre trial in women with a diagnosis of polycystic ovary syndrome (PCOS), obesity (BMI ≥ 30 kg/m2), and oligomenorrhoea or amenorrhoea (25). Participants were randomised to either metabolic bariatric surgery (Vertical Sleeve Gastrectomy) or medical care. The primary outcome was the number of biochemically confirmed ovulatory events over 52 weeks, where it was shown that metabolic bariatric surgery was more effective than medical care for the induction of spontaneous ovulation in women with PCOS, obesity, and oligomenorrhoea or amenorrhoea. Secondary outcomes were reproductive, metabolic, and quality-of-life outcomes over this period. Participants also had subcutaneous adipose tissue biopsies collected through an abdominal wall punch biopsy at baseline, 6, and 12 months following the intervention. The first two timepoints were analysed in this study (IRAS 269196).

Animals - All animal procedures undertaken were approved by the British Home Office under the UK Animal (Scientific Procedures) Act 1986 with approval from the local ethical committee (Animal Welfare and Ethics Review Board, AWERB), at the Central Biological Services (CBS) unit at the Hammersmith Campus of Imperial College London.

Adult male C57BL/6J mice (Envigo, Huntingdon U.K.) were maintained under controlled temperature (21-23°C) and light (12:12 hr light-dark schedule, lights on at 0700). The animals were fed either PMI Nutrition International Certified Rodent Chow No. 5CR4 (Research Diet, New Brunswick, NJ) or 58 kcal% Fat and 18.4% Sucrose diet (D12331, Research Diet, New Brunswick, NJ) to induce obesity and diabetes, for twelve weeks and ad libitum. Animals were exposed to liquid diet (20% dextrose) three days prior to surgery and remained on this diet for up to four days post operatively. Following this, mice were returned to either PMI or high fat/high sucrose diet. All mice were divided in two groups, VSG (n=4-5) and sham (n=4-5), and were euthanized, and tissues harvested, twelve weeks after surgery. Liver and adipose tissue biopsies were removed from all mice at twelve weeks following sham or VSG surgery in the fed state, and were either snap frozen in -80°C, fixed in formalin, or both. 11βHSD1^−/-^ mice lacking exon one, were generated by CRISPR/Cas9-mediated recombination, as previously described (26) on a C57BL/6J background. Heterozygous animals were inter-crossed to generate wild-type, heterozygous and homozygous littermates. Male mice were initially used as a continuation of our previous study (26) to allow direct comparison of results. Sample size calculation was performed with blood glucose as a primary outcome. Randomisation was performed according to pre-treatment body weight.

Vertical Sleeve Gastrectomy - Anaesthesia was induced and maintained with isoflurane (1.5-2%). A laparotomy incision was made, and the stomach was isolated outside the abdominal cavity. A simple continuous pattern of suture extending through the gastric wall and along both gastric walls was placed to ensure the main blood vessels were contained. Approximately 60% of the stomach was removed, leaving a tubular remnant. The edges of the stomach were inverted and closed by placing two serosae-only sutures, using the Lembert pattern. The initial full-thickness suture was subsequently removed. Sham surgeries were performed by isolating the stomach and performing a 1 mm gastrotomy on the gastric wall of the fundus. All animals received a five-day course of SC antibiotic injections (Ciprofloxacin 0.1mg/kg).

Glucose Tolerance Tests - Mice were fasted overnight (total 16 h) and given free access to water. At 0900, glucose (3 g/kg body weight) was administered via intraperitoneal injection. Blood was sampled from the tail vein at 0, 5, 15, 30, 60 and 90 min. after glucose administration. Blood glucose was measured with an automatic glucometer (Accuchek; Roche, Burgess Hill, UK).

Insulin tolerance tests - Mice were fasted for 8□h (08:00) and given free access to water. At 15:00, human insulin (Actrapid, Novo Nordisk) (1.5□U/kg body weight) was administered via intraperitoneal injection. Blood was sampled as described in Glucose Tolerance tests.

RNA isolation, cDNA synthesis and quantitative polymerase chain reaction (qPCR): RNA was extracted from cell and tissue samples by resuspending them in TRIzol reagent (Ambion, Life Technologies, Cat #349907) and chloroform (Sigma-Aldrich, Cat# SHBL1580). Total RNA isolation was conducted using the TRIzol™ Plus RNA Purification Kit (Applied Biosystems, Thermofisher) according to the manufacturer’s protocol. The High-Capacity cDNA Reverse Transcription kit (Applied Biosystems, Thermofisher) was used for cDNA synthesis. Master mix for qPCR was made using SYBR green (Applied Biosystems, ThermoFisher), RNase Samples were analysed using 7500 Fast Real-Time PCR system (Applied Biosystems, Serial #275010654). All data were normalised against β-actin as the housekeeping gene and all ΔCt values were calculated using the average from dual replicates. The analytical method was using 2-ΔΔCt. The primers used are shown in Table 1.

**Table 1:**
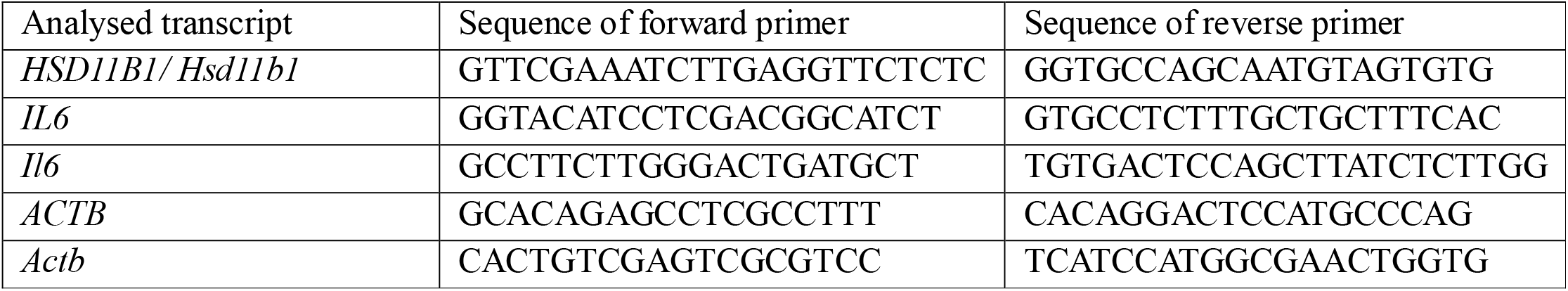
Primers used for the quantitative detection of transcripts normalized to β-actin.

Enzyme-linked immunosorbent assays - Blood plasma concentrations of IL6 (Proteintech) were measured using their rgespective assay kits and processed according to manufacturer’s protocol. Standard curves were interpolated by 5-parameter logistic curves. All quantified values between sensitivity limits and lowest standard points were included in the analysis. All values below quantification limits were also included in the analysis as sensitivity or ‘half-minimum’ values.

Cell culture and treatments - Cells were cultured with the aim of measuring 11βHSD1 expression in response to IL-6 cytokine treatment. HepG2 cells were purchased from ATCC (Manassas, VA) and were cultured with DMEM supplemented with 10% Foetal Bovine Serum, 1% Non-Essential Amino Acids and 1% Penicillin/Streptomycin (Invitrogen). Mycoplasma-free certificate was provided by the company. Cells were seeded at a density of□×□10^5^ per well and treated for 72 hrs with glucose (30mM) and IL-6 (1nM) (Abcam, USA). The medium was replaced every 24 hrs. A total of 4–10 different replicates were used for each condition.

Statistical Analysis – Each single patients was used as a unit of analysis (n=5). Each single animal was used as a unit of analysis (n=4 animals per group). Each cell culture experiment was repeated at least 3 times. Data were analysed using GraphPad Prism 10 software. Comparisons between two groups were carried out using Mann-Whitney test. Group comparisons were analysed using a two-way ANOVA, given lack of missing measurements, with Geisser-Greenhouse correction, and Sidak’s multiple comparison test. The model included main effects and interaction, to be able to evaluate group differences at individual timepoints. All statistical tests were 2-tailed, and significance was set at alpha = 0.05. Errors signify ± Median with interquartile range.

Data and resource availability - All data generated during this study are included in the published article

## Results

### Downregulation of 11βHSD1 and IL6 in subcutaneous adipose tissue following VSG in humans

Five female participants from the BAMBINI clinical trial underwent VSG (Table 2). Significant weight loss was observed (111.7 ± 6.9 kg at baseline vs 82.8 ± 9.3 kg at 6 months; 25.9% reduction, Fig. 1a), alongside an improvement in insulin resistance, reflected by reduced HOMA-IR (3.4 ± 1.5 vs 1.5 ± 0.6; 55.8% reduction, Fig. 1b). Gene expression analysis of subcutaneous adipose tissue revealed a significant reduction in the cortisol-activating enzyme 11βHSD1 at 6 months post-VSG (p < 0.01; Fig. 1c). Similarly, expression of the pro-inflammatory cytokine IL6 in adipose tissue was significantly decreased post-operatively (p < 0.1; Fig. 1d).

**Table 2.** Demographics table of participants.

| Ethnicity | Age | BMI |
| --- | --- | --- |
| British East Asian | 44.4 | 35.0 |
| White | 41.4 | 25.0 |
| White | 31.0 | 42.8 |
| White | 23.0 | 37.5 |
| White | 29.0 | 46.1 |
| Average | 33.8 | 37.3 |

**Figure 1.**
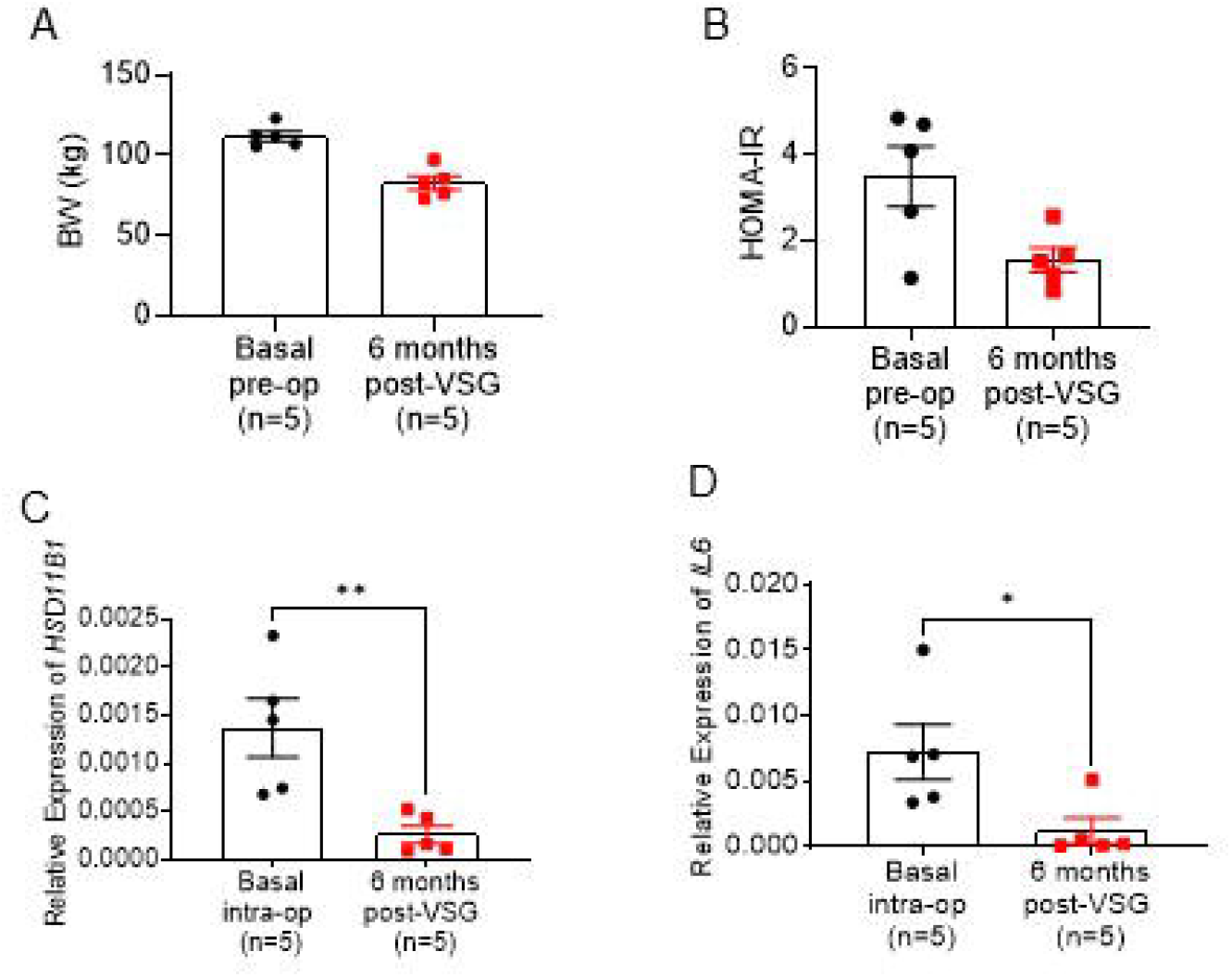
A. Body weight of patients at basal and 6 months post Vertical Sleeve Gastrectomy (VSG) B. HOMA-IR values of patients C. Gene expression of HSD11B1 and D. Interleukin 6 (IL6) relative to β-actin (ACTB), in subcutaneous adipose tissue biopsies collected intra-operatively and 6 months post VSG. (n=8/group) *p<0.05, **p<0.01

### VSG improves glucose tolerance independently of weight loss in lean mice

To determine whether these effects were driven by weight loss, VSG or sham surgery was performed in lean mice. Subcutaneous adipose tissue biopsies were collected intra-operatively and 4 weeks post-surgery. As expected in lean animals, no significant differences in body weight were observed between baseline and post-operative measurements (Fig. 2a). Despite the absence of weight loss, VSG-treated mice exhibited improved glucose tolerance during an oral glucose tolerance test (OGTT), with reduced glucose excursion compared to controls (AUC: 938.3 ± 135.6 vs 798.6 ± 126.3 mmol/L; Fig. 2b). Consistent with the human data, subcutaneous adipose tissue showed significant downregulation of both 11βHSD1 (p<0.001) and IL6 (p<0.01) gene expression following VSG (Fig. 2c,d), indicating that these changes are independent of weight loss.

**Figure 2.**
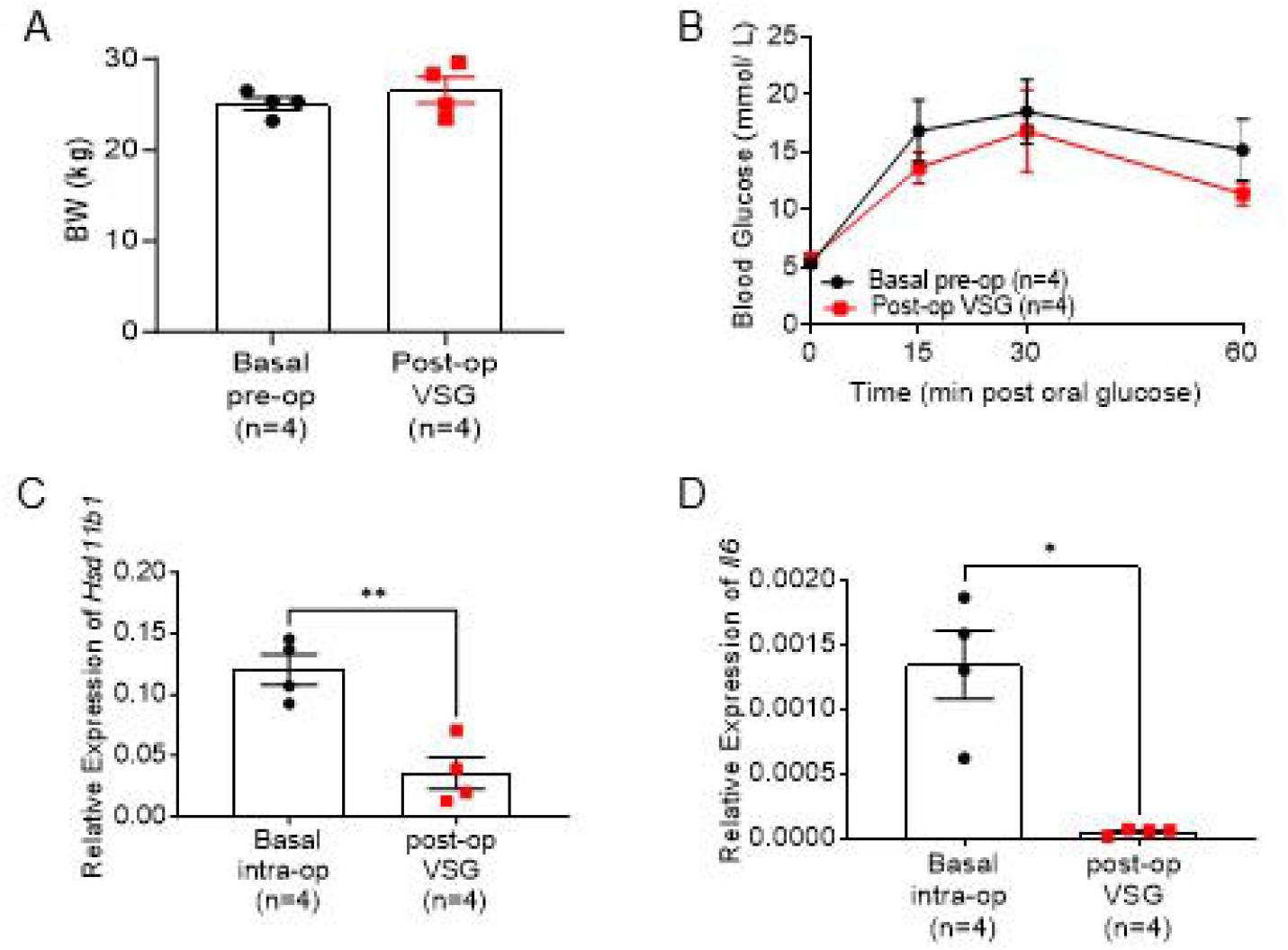
A. Body weight of lean mice at basal and 4 weeks post Vertical Sleeve Gastrectomy (VSG) B. Oral glucose tolerance test (3g/kg) following overnight fasting C. Gene expression of Hsd11b1 and D. Interleucin 6 (Il6) relative to β-actin (Actb), in subcutaneous adipose tissue biopsies collected intra-operatively and 4 weeks post VSG. (n=4/group) *p<0.05, ***p<0.0001

### Deletion of 11βHSD1 mimics the glucose-lowering effects of VSG in HFD mice

To investigate whether reduced 11β-HSD1 contributes to the metabolic benefits of VSG, we utilised global 11βHSD1 knockout mice alongside VSG and sham-operated C57BL/6J mice following 12 weeks of high-fat diet (HFD) (24). Unexpectedly, 11βHSD1 knockout mice exhibited greater weight gain compared to both sham and VSG groups (AUC: 161.4 ± 6.1 g vs 128.6 ± 3.2 g vs 110.5 ± 4.3 g; Fig. 3a). However, despite increased body weight, knockout mice displayed significantly improved glucose tolerance compared to sham controls, approaching the glycaemic profile observed in VSG-treated mice (AUC: 948.6 ± 73.1 vs 1196 ± 35.9 vs 835.5 ± 41.0 mmol/L; Fig. 3b). This improvement was associated with enhanced insulin secretion during the intraperitoneal glucose tolerance test (Fig. 3c). No significant differences were observed between groups during the insulin tolerance test (Fig. 3d), suggesting that insulin sensitivity was not markedly altered.

**Figure 3.**
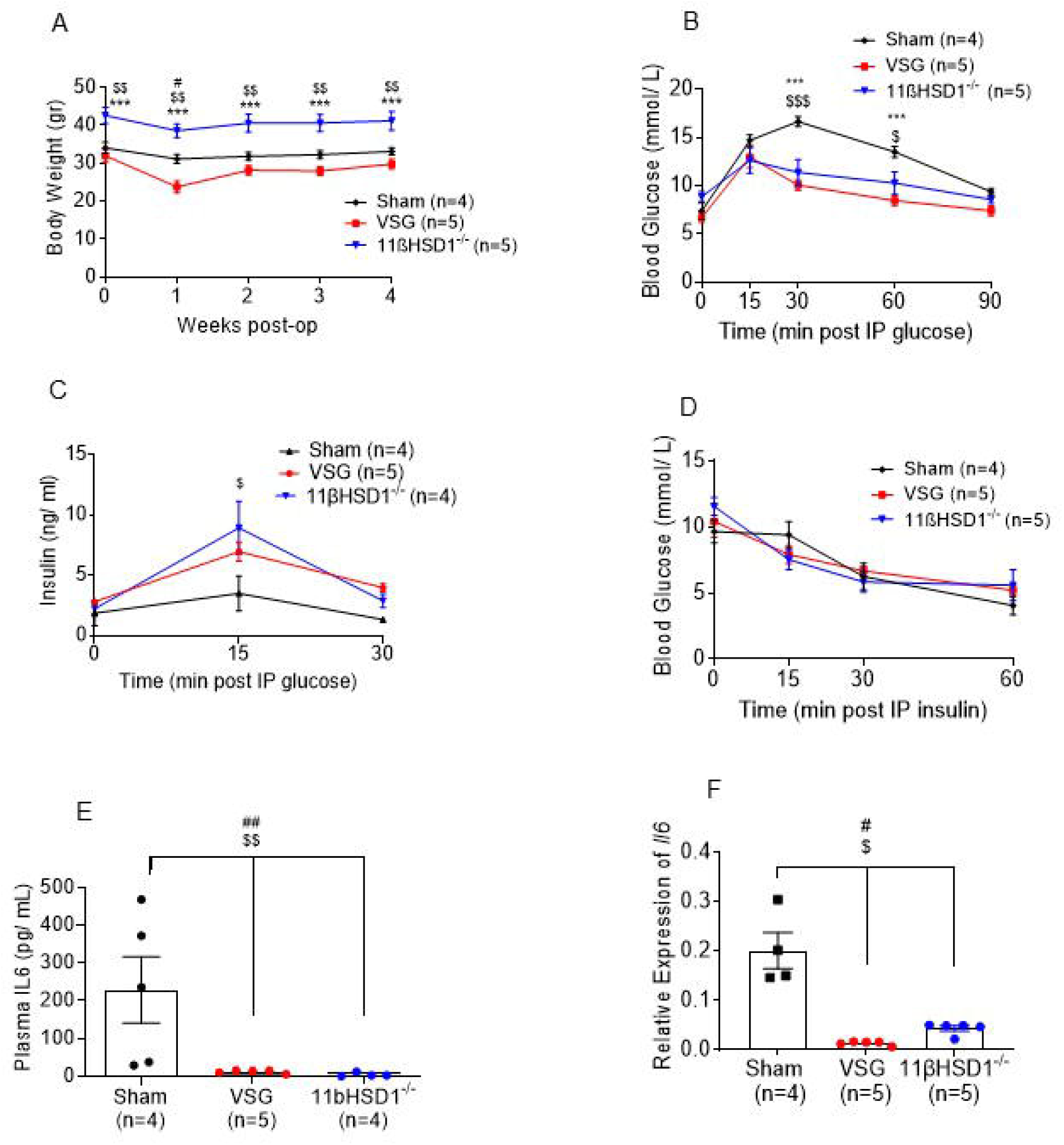
A. Body weight of high fat diet (HFD) wild type or 11βHSD1 knockout mice at basal and up to 4 weeks post Vertical Sleeve Gastrectomy (VSG) or sham surgery B. Intraperitoneal glucose tolerance test (1g/kg) following 6hr fasting C. Insulin secretion in plasma following 3g/kg intraperitoneal glucose injection D. Intraperitoneal insulin tolerance test (1.5IU/kg) following 6hr of fasting E. Plasma IL6 concentration F. Il6 relative to β-actin (Actb), in subcutaneous adipose tissue biopsies. All measurements took place 3-4 weeks post VSG/sham. (n=4-5/group) VSG vs. Sham #p<0.05, VSG vs. 11βHSD1 ***p<0.001, Sham vs. 11βHSD1 $ p <0.05, $$ p<0.001

### Reduced IL6 levels in both VSG-treated and 11βHSD1 knockout mice

Both plasma IL6 concentration and tissue gene expression were assessed across all groups. Both VSG-treated and 11βHSD1 knockout mice exhibited significantly reduced circulating IL6 levels compared to sham controls (p < 0.001; Fig. 3e). This reduction was mirrored in subcutaneous adipose tissue, where IL6 gene expression was similarly suppressed (p<0.01) in both groups (Fig. 3f).

### IL6 upregulates HSD11B1 expression in human hepatocytes

To further investigate the relationship between IL6 and 11βHSD1, human HepG2 hepatocytes were treated with 1nM IL6 for 72 hours.

Given that 11βHSD1 is most highly expressed in the liver, this model was selected to assess regulatory interactions. IL6 treatment resulted in a marked upregulation of HSD11B1 gene expression, with an approximately 18-fold increase compared to untreated controls (Fig. 4).

**Figure 4.**
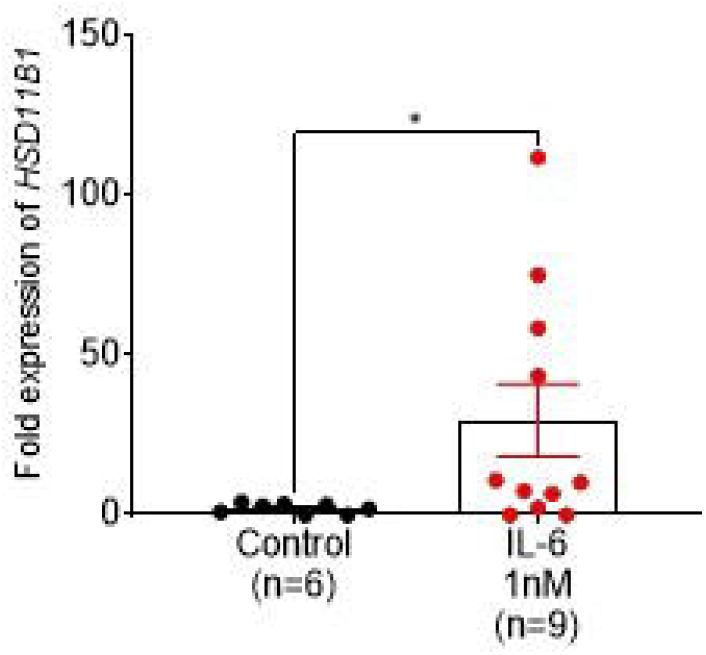
Relative expression of HSD11B1 in human hepatocyte cells (HepG2) treated for 72hrs with IL6. The culture medium was changed every 24 hrs.

## Discussion

In this study, we investigated the effects of VSG on glucocorticoid metabolism and inflammation in both humans and mice, with a particular focus on the role of 11βHSD1. We demonstrate that VSG is associated with a consistent downregulation of 11βHSD1 and IL6 expression in adipose tissue, alongside improved glycaemic control. Importantly, genetic deletion of 11βHSD1 in mice recapitulated key aspects of the glucose-lowering phenotype observed following VSG, supporting a functional role for this enzyme in mediating metabolic improvements. Furthermore, we provide evidence for a bidirectional relationship between inflammation and glucocorticoid metabolism, as IL6 stimulation upregulated 11βHSD1 expression in human hepatocytes.

Type 2 diabetes represents a major and growing clinical burden, characterized by impaired glucose homeostasis, insulin resistance, chronic low-grade inflammation, and dysregulated glucocorticoid metabolism (16, 27-29). While metabolic bariatric procedures such as VSG were initially developed for weight reduction, they are now recognized as highly effective metabolic interventions that improve glycemic control and insulin sensitivity (3-6). Notably, accumulating evidence indicates that these benefits cannot be fully explained by weight loss alone (24). Proposed mechanisms include altered gut hormone signalling, microbiome changes, and reduced systemic inflammation (8, 9, 30, 31). Our previous work also identified glucocorticoid regulation -specifically reduced 11βHSD1 expression as a potential contributor (24). Despite a known crosstalk between inflammatory pathways and the HPA axis (32, 33), the interplay between glucocorticoid metabolism and inflammation in this context has remained insufficiently defined.

Here, we show that VSG improves glucose tolerance in lean mice in the absence of significant weight loss, supporting the concept of weight-independent metabolic effects. While some studies attribute improvements in insulin sensitivity primarily to weight loss or energy deficit (34) (40-42), our findings align with evidence demonstrating rapid glycaemic improvements following metabolic bariatric surgery that precede substantial weight reduction. Previous work has largely focused on pancreatic β-cell function, insulin sensitivity, and incretin biology (35) (36, 37), whereas the contribution of glucocorticoid metabolism and inflammation has received comparatively little attention.

Our data further demonstrate that global deletion of 11βHSD1 improves glucose tolerance in HFD-fed mice, despite increased body weight. This finding is consistent with previous studies showing that Hsd11b1 knockout mice exhibit improved glycaemic control and enhanced insulin secretion, even in the context of obesity (38, 39) (40, 41). The observation that knockout mice partially phenocopy the metabolic effects of VSG suggests that suppression of 11βHSD1 contributes to, but does not fully account for, the benefits of surgery, indicating a multifactorial mechanism.

In humans, VSG resulted in significant weight loss, improved insulin resistance, and reduced expression of both 11βHSD1 and IL6 in subcutaneous adipose tissue. These findings are consistent with previous reports linking weight loss to reduced local cortisol regeneration and improved adipose tissue function (42, 43) (44) (4, 45) (46, 47). The concomitant decrease in IL6 further supports a reduction in adipose tissue inflammation, a key driver of insulin resistance (44) (48). While weight loss itself is associated with decreased IL6 levels (49), our parallel findings in lean mice indicate that these anti-inflammatory and glucocorticoid-related effects can occur independently of weight reduction.

Taken together, our results support a model in which VSG induces coordinated changes in glucocorticoid metabolism and inflammatory signalling. Both VSG and 11βHSD1 deletion were associated with reduced IL6 levels in plasma and adipose tissue, suggesting that decreased local cortisol activation may contribute to an anti-inflammatory state. Previous studies have similarly shown that inhibition or deletion of 11βHSD1 attenuates inflammatory responses in various tissues (50) (51), while VSG has been associated with reduced systemic and tissue-specific inflammation (16) (49).

Our *in vitro* findings extend this relationship by demonstrating that IL6 directly upregulates 11βHSD1 expression in human hepatocytes, indicating a bidirectional interaction. While glucocorticoids are classically anti-inflammatory, inflammatory stimuli have been shown to enhance local glucocorticoid activation through upregulation of 11βHSD1(52, 53). This process is thought to involve transcriptional regulation via C/EBPβ and NFκB pathways (53). Together, these findings suggest the existence of a feedback loop in which inflammation drives glucocorticoid activation, which in turn modulates inflammatory responses.

This study is strengthened by its translational approach, integrating human, mouse, and *in vitro* data to investigate the relationship between glucocorticoid metabolism, inflammation, and glucose homeostasis following VSG. The use of 11βHSD1 knockout mice provided mechanistic insight and supported a functional role for 11βHSD1 in mediating metabolic improvements. Furthermore, the observation of improved glucose tolerance in lean mice supports weight-independent effects of VSG. However, several limitations should be acknowledged. The use of global knockout mice limits tissue-specific interpretation and may involve developmental compensatory mechanisms. In addition, the inflammatory analysis focused primarily on IL6. Finally, although associations between reduced 11βHSD1 expression and improved metabolic outcomes were observed in humans, causality could not be definitively established, and further studies using pharmacological inhibition and tissue-specific models are needed.

Overall, our study identifies 11βHSD1 as a key node linking glucocorticoid metabolism, inflammation, and glucose homeostasis in the context of VSG. The parallel effects observed with surgical intervention and genetic deletion highlight the therapeutic potential of targeting this pathway. Further studies are needed to determine whether pharmacological inhibition of 11βHSD1 can replicate the metabolic benefits of metabolic bariatric surgery in humans.

## Author contribution

CRediT: Conceptualization: EA. Formal analysis: EA, SS, SL, IDD, WW, HLPT, AR. Funding acquisition: EA, ADM. Methodology: EA, AR, ADM. Project administration: EA, SS, BJ, ADM. Visualization: EA. Writing – original draft: SS, SL, EA. Writing – review & editing: EA, ADM. EA is the guarantor of this work and, as such, had full access to all the data in the study and takes responsibility for the integrity of the data and the accuracy of the data analysis.

